# Puckered in pioneer neurons coordinates the motor activity of the *Drosophila* embryo

**DOI:** 10.1101/092486

**Authors:** Katerina Karkali, Samuel W. Vernon, Richard A. Baines, George Panayotou, Enrique Martín-Blanco

## Abstract

Central Nervous System (CNS) organogenesis is a complex process that obeys precise architectural rules. The impact that nervous system architecture may have on its functionality remains, however, relatively unexplored. To clarify this problem, we analyzed the development of the Drosophila embryonic Ventral Nerve Cord (VNC). VNC morphogenesis requires the tight control of Jun kinase (JNK) signaling, exerted in part via a negative feedback loop mediated by the dual specificity phosphatase Puckered, in a subset of pioneer neurons. Here we show that the JNK pathway autonomously regulates neuronal electrophysiological properties without affecting synaptic vesicle transport. Manipulating, during early embryogenesis, JNK signaling activity in pioneer neurons, directly influences their function as ‘organizers’ of VNC architecture and, moreover, uncovers a role in the coordination of the embryonic motor circuitry that is required for hatching. Together, our data reveal critical links, mediated by the control of the JNK signaling cascade by Puckered, between the structural organization of the VNC and its functional optimization.

## INTRODUCTION

Nervous system architecture is the outcome of neuro-glial interactions and axon guidance decisions. Yet, how the nervous system achieves a mechanically balanced organization while functional neuronal connectivity is established throughout morphogenesis is poorly understood. Several lines of evidence suggest a direct relationship between the 3D architectural organization of the nervous system and its functionality. First, the spatial distribution of functional circuits in the cortex of vertebrates extensively overlaps with large-scale microstructure, connectivity, and gene expression patterns ^1^. Second, in pursuit of functional efficiency, neural networks likely minimize their structural cost following the wiring optimization principle postulated by Ramón y Cajal ^2^. These costs likely arise from metabolic requirements, signal delay and attenuation, and complex guidance processes ^3^ that increase with interneuronal distance ^4^. Lastly, the spatial organization of cortical areas appears subjected to evolutionary selection ^5^.

We have recently shown that the acquisition of the *Drosophila* embryonic Ventral Nerve Cord (VNC) specific 3D neural architecture is directed by the mechanical coordination of neurons and glia ^6^. Further, we have also uncovered that a precise modulation of the activity of the JNK signaling cascade, in a subset of early specified neurons, is required for proper VNC architectural organization and its condensation (Karkali et al, co-submitted). We now determine whether accuracy in nervous system spatial organization/topography is essential for functionality. In particular, by employing the VNC as a model system, we asked if JNK-mediated modulation of VNC architecture is essential for the emergence of the coordinated embryonic motor circuit activity that mediates larval hatching ^7^

The best-known role of the JNK signaling cascade in the nervous system is to coordinate the induction of protective genes in response to oxidative (ROS) challenges ^8^ and axonal regeneration ^9^. Yet, JNK has been found to mediate multiple functions in the CNS both in *Drosophila* and mice, including axonal growth ^10^; axonal organelle (mitochondrial) and synaptic protein transport ^8, 11, 12, 13, 14^; synaptic growth and synaptic strength ^15^; dendrite pruning ^16^; self-renewal of neuronal precursors ^17^; neuronal migration ^18^; glia remodeling ^19^ and neuronal neuroplasticity (reviewed in ^20^). Further, and importantly, it is well established that some of the effects of JNK activation in the CNS do not proceed through the regulation of the transcription machinery (AP1), and likely occur via phosphorylating cytoplasmic targets in the soma ^8, 13^. In this scenario, we hypothesize that the JNK pathway may have a regulatory role in the optimization of neural function, associated with the structural organization of the VNC. During VNC condensation, *Drosophila* embryos develop sporadic muscle contractions that initially do not correlate with sensory input ^21, 22^. Yet, over time, these contractions evolve into coordinated peristaltic patterns, dependent on a precise coordination of the contractile activity of the embryonic muscles, bilaterally and along the antero-posterior (AP) axis ^7^. Here, we uncover essential functions for the JNK pathway in a subset of pioneer neurons enabling the emergence of coordinated motor activity. Early in embryogenesis, a tight control of JNK signaling is necessary, to attain an accurate axonal scaffold organization and native dendrites and axons conformation. Moreover, it is essential for the development of the complex pattern of muscle activities leading to hatching behavior. At later embryonic stages, JNK activity seemingly fine-tunes cell soma location and dendritic morphology and number. Likewise, axonal transport of synaptic vesicles appears to be unaffected by JNK activity, but is critical, upon synaptic input in pioneer motoneurons, for reaching the right Resting Membrane Potential (RMP) to support action potential firing. Combined, our data uncover fundamental structural and physiological roles for the JNK signaling pathway in pioneer neurons linked to embryonic motor pattern optimization.

## RESULTS

### The level of JNK activity in pioneer neurons is essential for embryonic motor coordination

Considering the roles of JNK signaling on promoting VNC architectural robustness (Karkali et al, co-submitted), we investigated the potential functional consequences of releasing the negative feedback control of JNK activity implemented by Puc. We first focused on motor circuits and evaluated late embryonic activity patterns ^7, 21^ (**Figure 1A** and **1B**) (see **Methods** and **Movie S1**). In wild type conditions, developing embryos execute forward and backward waves of peristalsis by coordinating bilaterally symmetrical muscle contractions ^7, 23^. We found that these movements matured through five stages (**Figure 1C**): they began with brief, isolated, unilateral twitches and sporadic head pinching (stage A). This initial phase was followed by a prolonged rest period, occasionally interrupted by uncoordinated single contractions, (B) and then by an active stage showing frequent incomplete and complete peristalsis (C). The active peristalsis stage precedes a short resting period (D), prior to the violent head movements and multiple complete peristaltic events (E) leading to the rupture of the vitelline membrane and larval hatching (see also **Figure S1** and **Movie S2**). In the complete absence of *puc* (*puc*^*E69*^), when embryos fail to condense their VNC (Karkali et al, co-submitted), the progression of motor patterns was interrupted, and embryos remained indefinitely in an immature stage (A) characterized by occasional unilateral twitches and incomplete peristalsis (**Figure 1D** and **Movie S3**). Muscle integrity and functionality were not affected.

**Figure 1.**
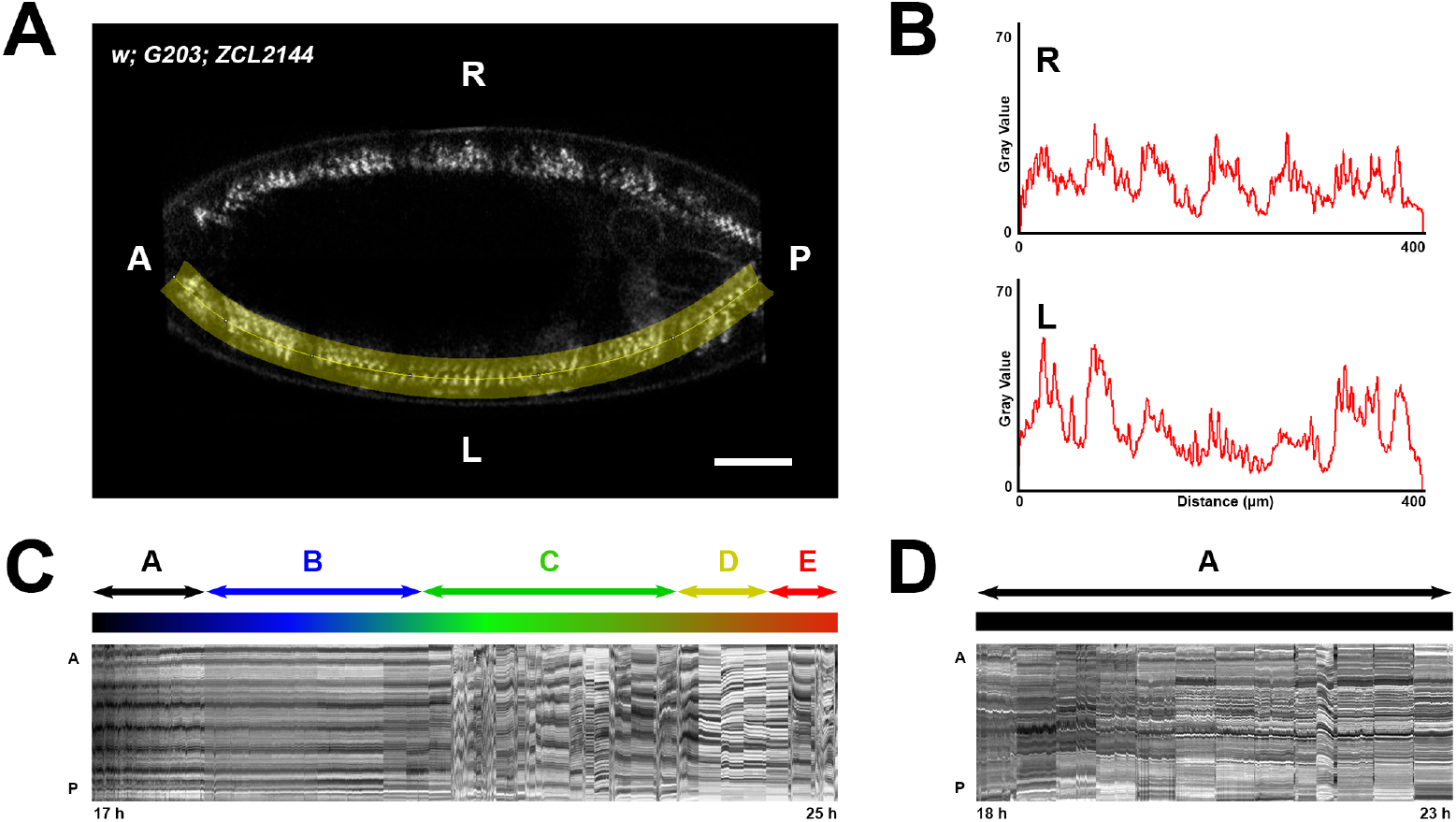
Defective motor coordination results in incomplete peristalsis in *puc*^*E69*^ embryos. **A)** Representative mid-plane image of a stage 17 embryo carrying two GFP protein traps expressed at muscle Z-lines (*w; G203; ZCL2144*) (see **Movie S1**). A selected segmented-line (highlighted in yellow) was employed to create kymographs spanning muscle units along the A/P axis (see below) at both sides of the embryo, right (R) and left (L). The evolution overtime of these kymographs was monitored live. Scale bar is 25 μm. **B)** Intensity Profile plots generated along the segmented-line selection in **A**. Gray values as a function of distance highlight the distribution of the segmental muscle units along the axis. Muscle contraction is detected by the differences in the distance between consecutive peaks at sequential time points. **C)** Kymograph displaying muscle profiles from 17 to 25 hours AEL for the left side of the embryo in **A**. Anterior (A) and posterior (P). Motor coordination maturation occurs in 5 stages of approximated stereotyped length (color coded): A, Intense unilateral twitching; B, general inactivity with occasional uncoordinated contractions; C, initiation of backward and forward peristalsis; D, pre-hatching inactivity period and E, complete peristalsis and specialized hatching head movements. **D)** Kymograph displaying muscle profiles from 18 to 23 hours AEL, when the embryo dies, for the left side of a *puc*^*E69*^ embryo (see **Movie S8**). Anterior (A) and posterior (P). Motor coordination fails and embryos do not progress beyond the unilateral contraction period (stage A).

The systemic failure of motor coordination observed in *puc* mutants could be a direct consequence of the increase of JNK activity levels in the CNS or the outcome of other non-defined non-autonomous events. To discard this last possibility, we mimicked *puc* loss of function in different subsets of *puc* expressing neurons such as the aCC, pCC and RP2 pioneers (RN2 Gal4) and the VUM midline neurons (MzVUM Gal4) by overexpressing a constitutively active form of the JNKK (Hep^CA)^). Importantly, interfering in JNK activity in these cells results in architectural defects and in VNC condensation arrest (Karkali et al, co-submitted). For RN2 (**Figure 2B** and **2C**) or MzVum cells (**Figure 2D** and **2E**), an excess of JNK activity resulted in a phenotype equivalent to that observed in *puc*^*E69*^ mutant embryos. Only occasional unilateral twitches, head pinches and infrequent incomplete peristalsis were observed. Consequently, the motor defects observed in *puc* appear to be a direct response to the loss of Puc activity in specific *puc*+ neurons. The misregulation of the JNK activity links both VNC structural and embryonic motor coordination defects.

**Figure 2.**
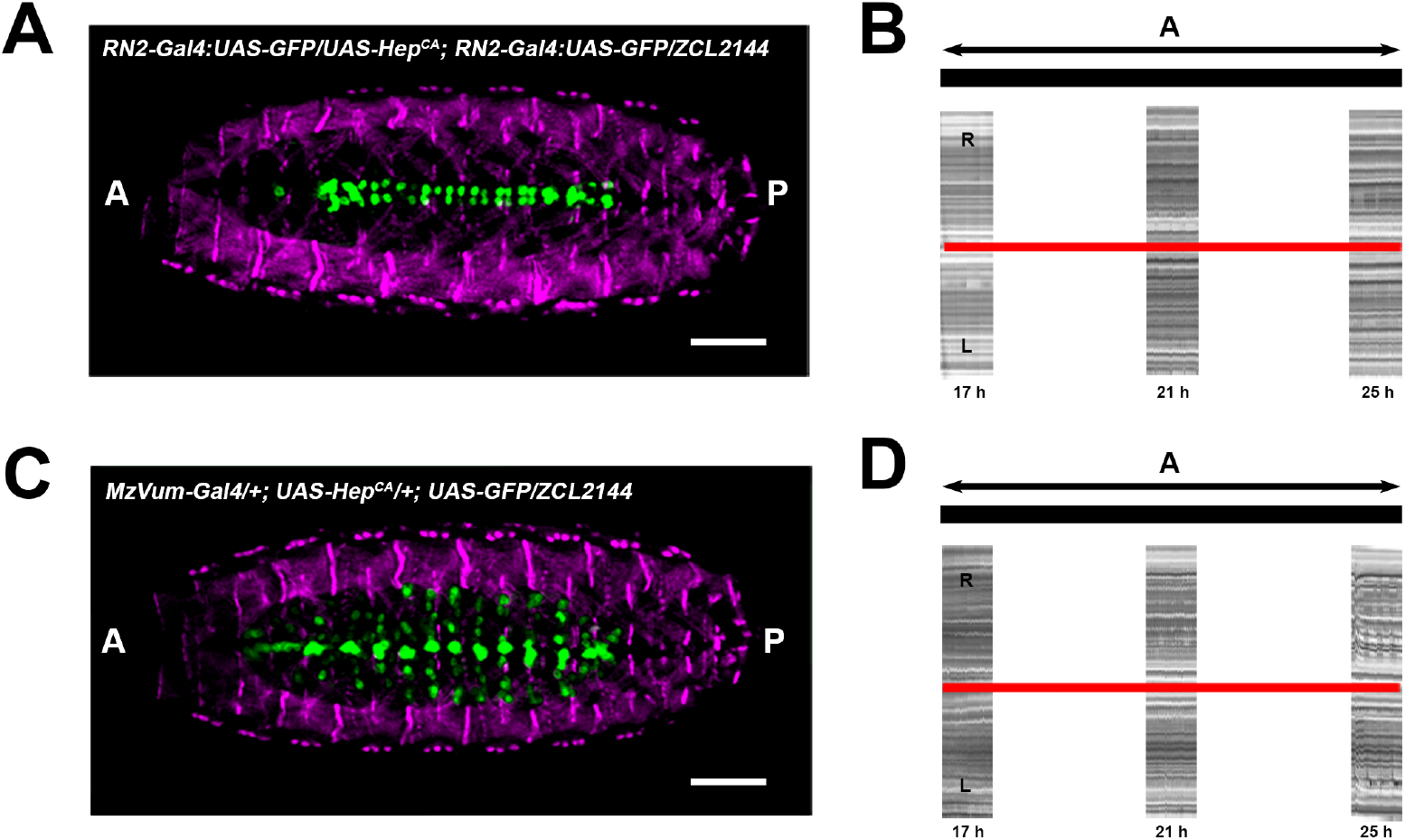
JNK activity modulates embryo motor coordination. **A** and **C** Ventral view of stage 17 embryos expressing GFP under the control of the RN2-Gal4 (**E**) and MzVum-Gal4 (**G**) lines co-expressing Hep^CA^ and carrying the muscle Z-lines specific GFP protein trap ZCL2144. Muscles are pseudo-colored in magenta and neurons in green. Scale bar is 25 μm. **B** and **D**) Kymographs (ZCL2144 marker) revealing occasional unilateral contractions in embryos co-expressing Hep^CA^ in the RN2-Gal4 (**F**) and MzVum-Gal4 (**H**) neurons. For each case, the kymographs correspond to three periods of 1 hour. These embryos never manage to coordinate muscle movements. Both sides, right (R) and left (L) are sh

### JNK signaling does not influence synaptic vesicle transport but modulates motoneuron firing

Would the loss of embryo motor synchrony after altering JNK activity in *puc*-expressing neurons be consequence of the aberrant architectural organization and inability of the VNC to condense? Some *puc*+ cells are motoneurons and at anomalous levels of JNK activity, their signaling capability might be impaired. This might result in defects in the embryo’s motor coordination independently of the VNC architectural organization. To evaluate this possibility, we first studied the axonal transport of synaptic vesicles (Synaptotagmin - Syt) and organelles (Mitochondria - Mito) in the intersegmental nerve [aCC and RP2 (RN2-Gal4) and in VUM motoneurons (MzVum-Gal4)] (see **Methods, Figures 3 and S2** and **Movie S4**). Both in RN2 and MzVum axons, synaptotagmin particles moved at a speed of around 0,7 μm/s with a tracking lifetime of 50 s per event. Mitochondria run at the same average speed but these movements were less persistent and remained on track 30 s on average. Particles densities (frequency of events) were equivalent for synaptotagmin and mitochondria but larger for aCC and RP2 (0,3 N/μm) than for VUMs (0,2 N/μm). No differences were observed between anterograde and retrograde transport. Next, we analyzed particle transport upon co-expression of Hep^CA^. Synaptotagmin vesicles kinematic behavior was largely unaffected (**Figure S2**), which indicates that microtubule-mediated transport remained fully functional. The density of traveling mitochondria, however, dropped in aCC and RP2 and increased in VUMs (**Figure 3**), yet sustaining their average lifetime. We noticed the presence of a subpopulation of fast moving mitochondria that could reach a speed of up to 3 μm/s. As for the wild type, we could not distinguish differences between anterograde and retrograde transport. In conclusion, transport of vesicles to synaptic terminals does not seem to be affected by the deregulation of JNK activity. Mitochondrial mobility, however, possibly as a result of altered energetic demands, became anomalous.

**Figure 3.**
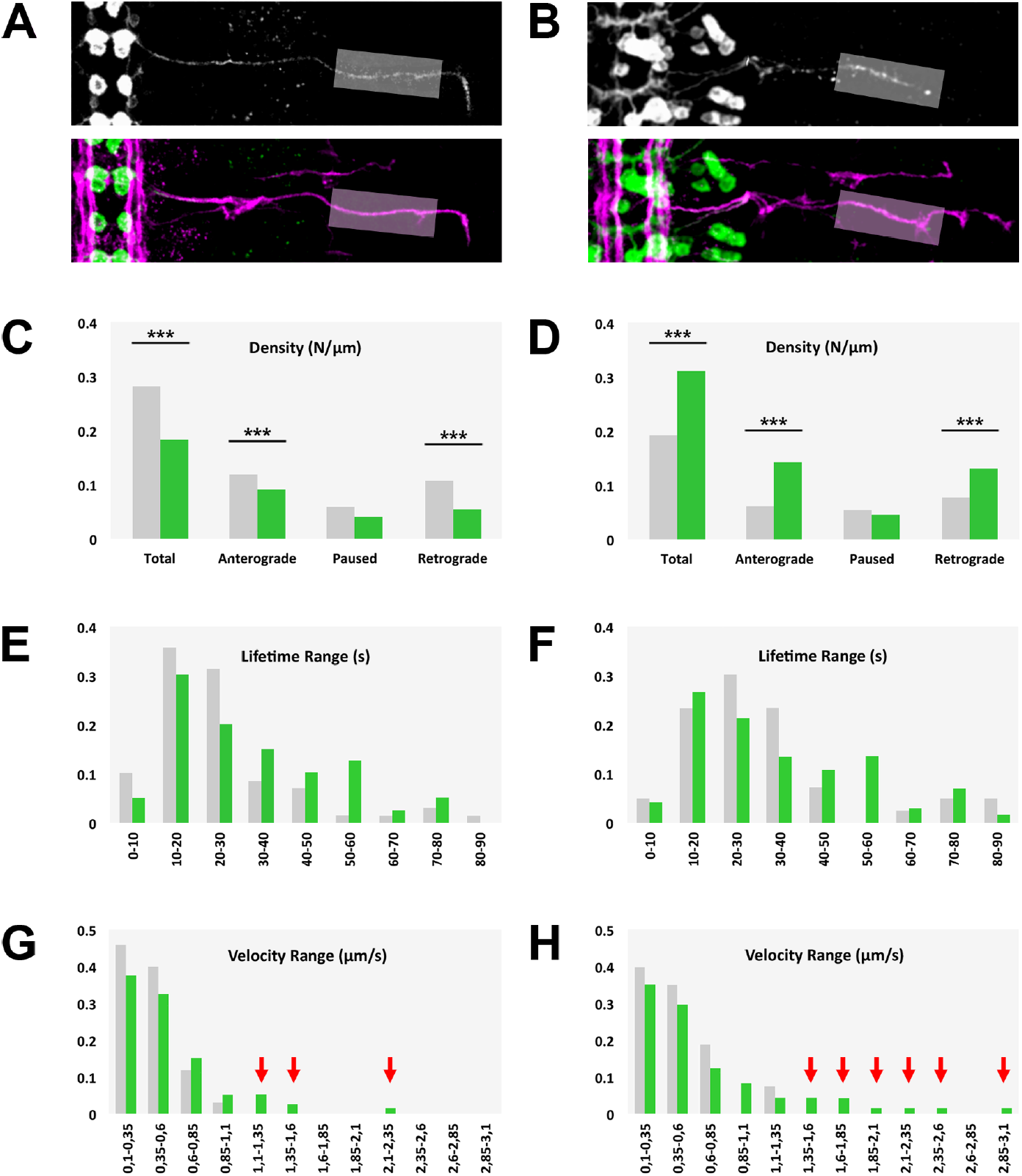
JNK activity affects mitochondrial axonal transport. **A** and **B**) Flat-prepped, stage 16 embryos expressing Mito-GFP under the control of the RN2-Gal4 (**A**) and MzVum-Gal4 (**B**) lines immunostained for GFP (gray scale - top panels and green - bottom panels) and Fas 2 (magenta - bottom panel). The grey rectangle marks the medial part of the ISN, where mitochondrial axonal transport was monitored (see **Movie S3**). Scale bar is 25 μm. **C** and **D**) Histograms showing mitochondrial density and directional motility in control (grey) versus Hep^CA^ expressing (green) aCC/ RP2 (**C**) and VUM motoneurons (**D**). Statistically significant differences (p<0.001) were observed between control and JNK overactive neurons. **E** and **F**) Lifetime range distribution of motile mitochondria in aCC/RP2 (grey) and Hep^CA^ expressing (green) embryos. **G** and **H**) Mean velocity range distribution comparison of motile mitochondria in aCC/RP2 (**G**) and VUM (**H**) motoneurons between control (grey) and JNK gain-of function (green) conditions. For both sets of motoneurons the distribution was shifted to higher velocity classes in JNK hyperactive neurons.

To directly address if the JNK activity influences motoneuron function, we undertook patch clamp recordings from the aCC and RP2 motoneurons, which receive identical cholinergic synaptic inputs ^24^. Neurons were voltage-clamped at −60mV to record endogenously produced cholinergic synaptic currents [Spontaneously Rhythmic Currents (SRCs)]. We found that expression of Hep^CA^ in RN2 + neurons did not influence SRC amplitude or frequency (**Figure 4A** and **4B**), indicating that the synaptic input to aCC/RP2 remained unaffected. However, current clamp recordings, in which no holding potential was applied (I=0), revealed that the motoneuron Resting Membrane Potential (RMP) was significantly increased and the membrane hyperpolarized (**Figure 4C** and **4D**). Therefore, the average number of action potentials fired per Excitatory Post Synaptic Potential (EPSP) was also notably reduced. We conclude that although the synaptic drive was normal, upon JNK hyperactivation, due to a more negative RMP, excitability, and thus endogenous firing, was considerably reduced in pioneer motoneurons.

**Figure 4.**
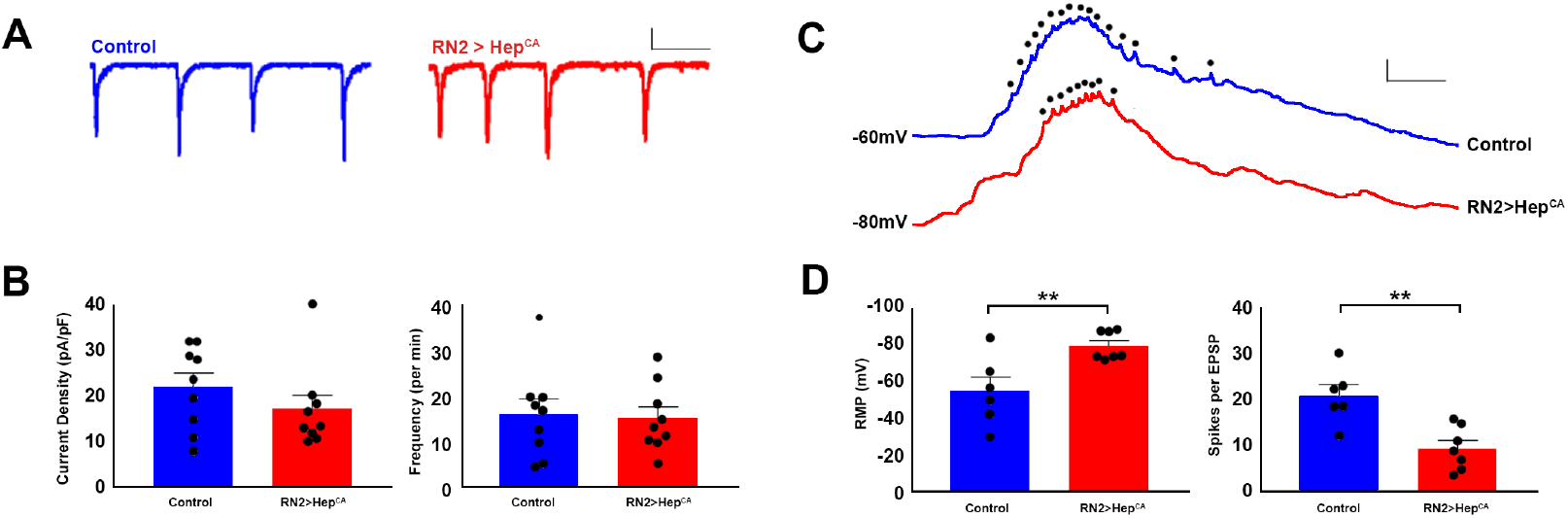
JNK activity modulates neuronal firing. **A)** Representative traces of endogenously produced cholinergic SRCs from aCC/RP2 motoneurons in late stage 17 embryos in control (*RN2-Gal4 / CyO*) and experimental lines (*RN2-Gal4 > UAS-Hep*^*CA*^). Scale Bar: 20 pA / 2s. **B)** No significant difference in SRC amplitude (21.7 ± 3.0 vs. 17.0 ± 3.1 pA / pF, control vs. *RN2-Gal4 > UAS-Hep*^*CA*^ respectively, P = 0.28) or frequency (16.3 ± 3.3 vs. 15.4 ± 2.4 per min, P = 0.83) was observed. **C)** Representative traces of endogenously produced cholinergic EPSPs from aCC/RP2 motoneurons in late stage 17 embryos in control (*RN2-Gal4 / CyO*) and experimental lines (*RN2-Gal4 > UAS-Hep*^*CA*^). Action potentials are indicated by dots. The amplitude of the action potentials is shunted because of the underlying depolarization. Scale Bar: 10 mV / 100ms. **D)** Motoneuron resting membrane potential (RMP) is significantly hyperpolarized in *RN2-Gal4 > UAS-Hep*^*CA*^ (−57.0 ± 7.6 vs. −81.5 ± 2.9 mV, control vs. *RN2-Gal4 > UAS-Hep*^*CA*^ respectively, P = 0.009). Consequently, the average number of action potentials fired per EPSP was also significantly reduced (20.6 ± 2.4 vs. 9.3 ± 1.7 spikes per EPSP, P = 0.003). All data points are shown on graphs, bars represent mean ± sem.

### JNK signaling influences the axodendritic organization of *puc*-expressing neurons

The JNK signaling pathway does not affect synaptic vesicle transport or neuron sensory competencies but modulates their firing response. Could this be related to potential alterations on pioneer neuron morphology and dendrite integrity? To explore this possibility, we examined, for structural defects, the morphology of *puc*-expressing pioneer neurons [RN2 (aCC, pCC and RP2)] upon Hep^CA^ overexpression.

aCC is a motoneuron whose axon gets incorporated into the intersegmental nerve (ISN) ^25^. Its dendritic arbor develops in a distinct region of the ISN myotopic dendritic domain and consists of 8 to 10 dendrites ^26^ of an average length of 4 μm. pCC is a medially-located interneuron whose axon extends ipsilateral, in an anterior direction ^27^. Finally, RP2 is a motoneuron, whose axon proceeds through the posterior root of the ISN ^25, 28^ and branches out between 7 and 9 dendrites, with an average length of 4 μm (**Figure 5A**).

**Figure 5.**
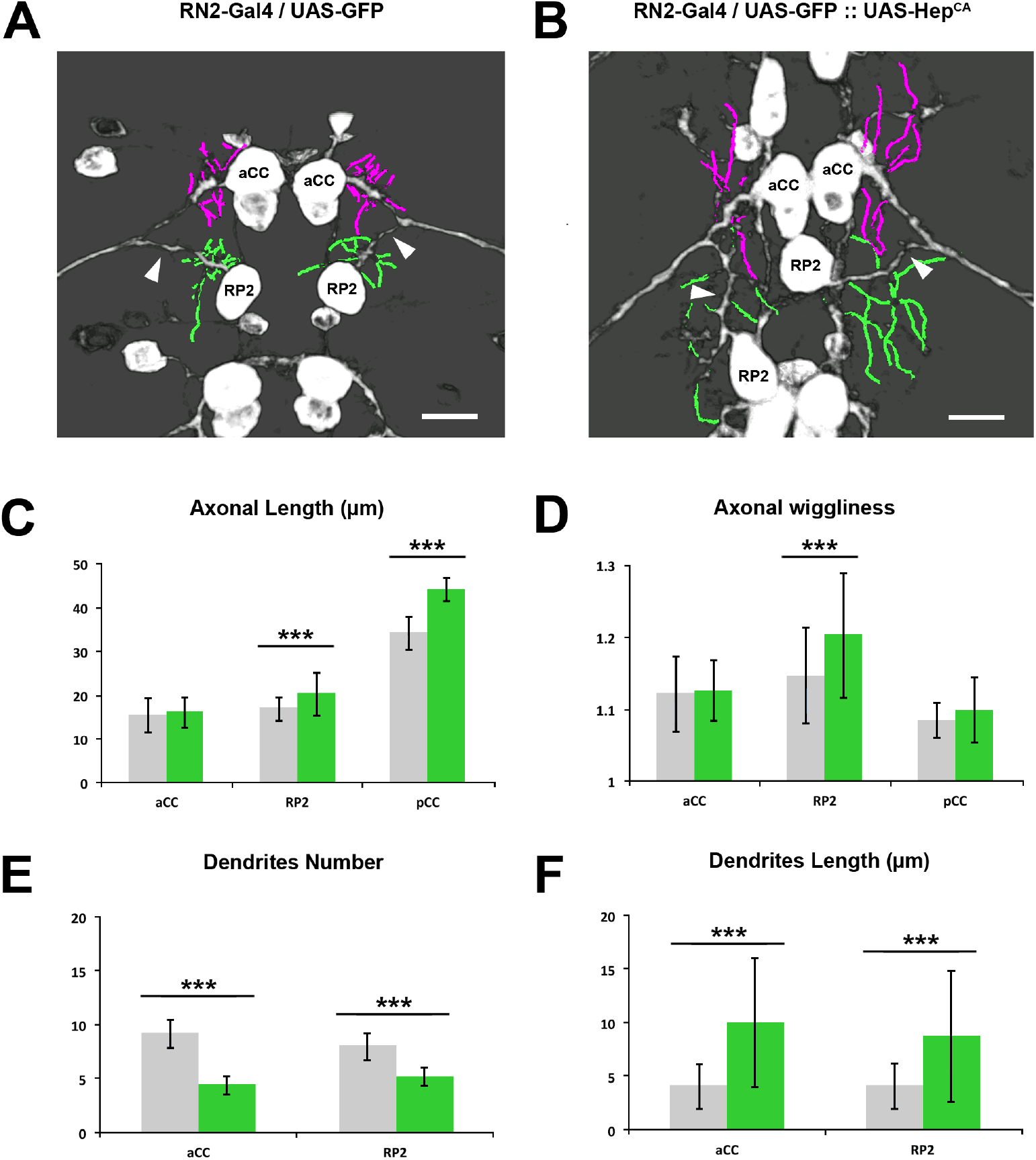
The axonal and dendritic landscape of pioneer neurons is altered upon JNK hyperactivation. **A** and **B**) Flat-prepped, stage 16 embryos expressing mCD8-GFP (**A**) and mCD8-GFP and Hep^CA^ (**B**) under the control of the RN2-Gal4 line immunostained for GFP. Traces of the aCC (magenta) and RP2 (green) proximal dendrites are over-imposed. Yellow arrowheads point to differences in RP2 axon morphology. Anterior is up. Scale bar is 10 μm. **C**) Histograms showing the average axonal length of the aCC, RP2 and pCC neurons. Statistically significant differences (p<0.001) in the axonal length of the RP2 and pCC neurons were detected between control and Hep^CA^ expressing neurons (n= 4 embryos). Control embryos are represented in grey and Hep^CA^ expressing embryos in green. **D**) Histograms showing the average axonal wiggliness of the aCC, RP2 and pCC neurons. RP2 axonal wiggliness was found significantly increased after Hep^CA^ expression (p<0.001, n= 4 embryos). **E** and **F**) Graphs representing the average Dendrite Number (**E**) and average Dendrite Length (**F**) of the aCC and RP2 neurons. In all cases significant differences were scored in the comparison of control and Hep^CA^ expressing embryos (p<0.001, n= 4 embryos).

As a consequence of the overexpression of Hep^CA^ in RN2+ neurons, the cell bodies of the aCC, pCC and RP2 were frequently misplaced and came together towards the midline (compare **Figures 5A** and **5B**). The appearance of their axons was also affected; they thickened (**Figure 5B**), increased in length (for pCC and RP2) (**Figure 5C**) and developed a distinctly different wiggly morphology (**Figure 5D**). JNK hyperactivation also affected the density (**Figure 5E**) and length (**Figure 5F** and **Movie S5**) of dendrites. Summarizing, increasing JNK activity in specific pioneer neurons leads to local structural and topographical disarrays that could seed for mistaken 3D arrangements and cumulative misrouting, and be related to their impaired functionality.

### Different temporal roles for the JNK pathway

To define if the inputs of the JNK pathway modulating the structural organization and functional optimization of the VNC are causally related or independent, we evaluated the effects of interfering in the JNK activity levels within the VNC at different times.

Programmed temperature shifts between 29 and 18ºC or *vice versa* (see **Methods**) allowed us to selectively overexpress Hep^CA^ early vs late during embryogenesis. The early-only (stage 10-13) hyperactivation of JNK in RN2+ neurons, before the initiation of neural activity ^29^, was associated with defects on the morphology of dendrites and axons, with axonal scaffold anomalies and with the collapse of RN2+ neurons towards the midline (compare **Figures 6A, B and C**). VNC condensation was also affected (**Figure 6E**). Furthermore, although these embryos occasionally hatched, they displayed strong muscle coordination defects with an almost complete absence of bilateral peristalsis (**Figure 6F** and **Movie S6**). Conversely, the late hyperactivation of the JNK pathway (stage 15-16) did not affect VNC condensation, neither motor coordination and peristalsis (**Figures 6E** and **6F** and **Movie S6**). However, it resulted in the misalignment of RN2+ neurons, which displayed scarce and long dendrites, and in the formation of heterogeneous axon bundles (**Figure 6D)**.

**Figure 6.**
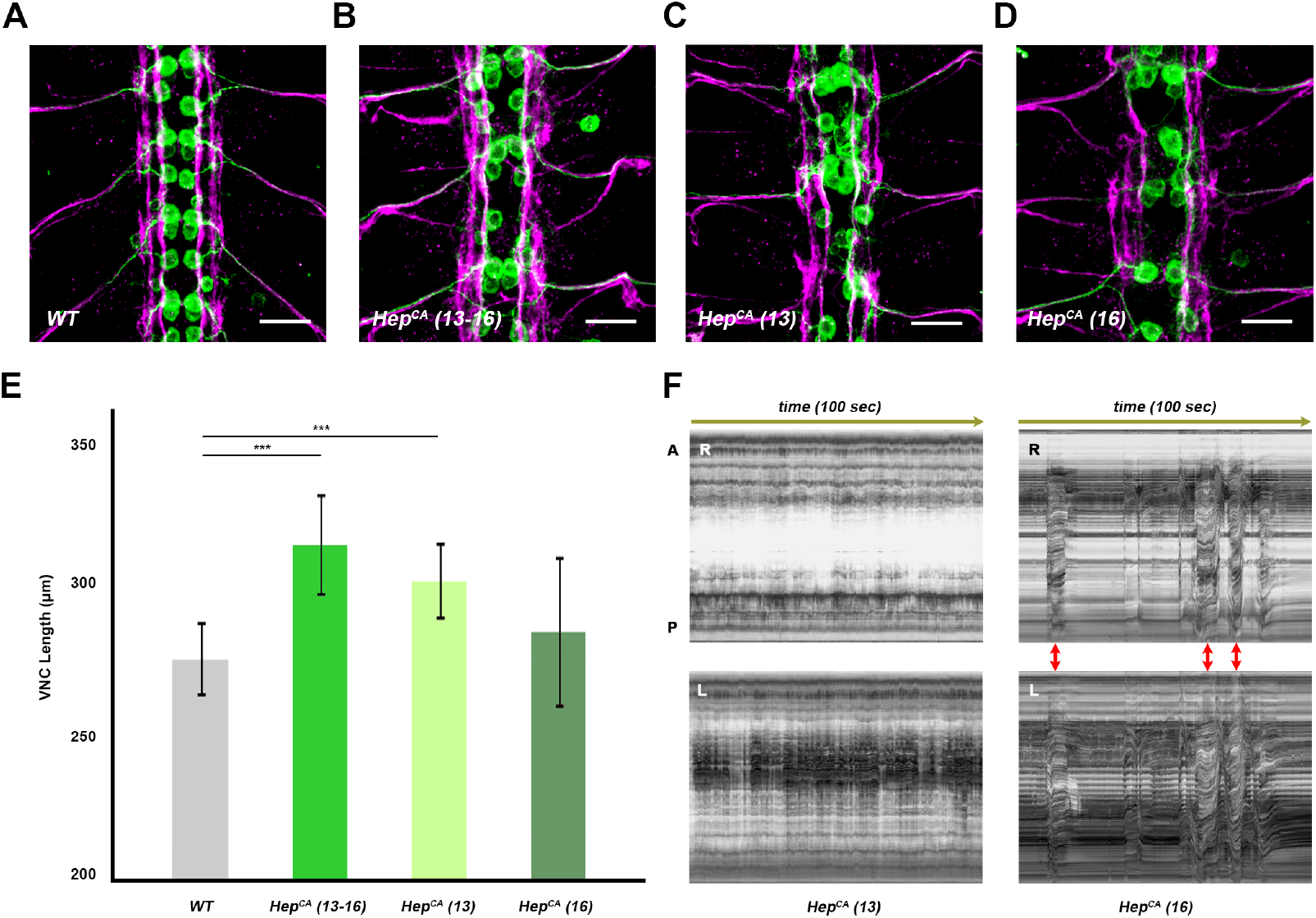
JNK activity modulates VNC condensation and embryo motor coordination. **A-D)** Fas 2 (magenta) and GFP (green) immunoreactivity of RN2-Gal4 embryos at late stage 16 subjected to different temperature shift conditions and expressing, from left to right: (**A**) GFP alone (WT) (maintained at 29ºC continuously from stage 10 to 16); or GFP in combination with a *Hep*^*CA*^ transgene; Hep^CA^ (10-16) (maintained at 29ºC continuously from stage 10 to 16) (**B**); Hep^CA^ (10-13) (kept at 29ºC between stages 10 and 13) (**C**) and Hep^CA^ (15-16) (kept at 29ºC during stages 15 and 16) (**D**). Maximum projection of ventral views across three VNC abdominal segments. Scale Bar is 10 μm. Overexpressing *Hep*^*CA*^ at early stages affects pioneer functions and provoke the collapse of neurons cell bodies into the midline. At late stages, dendrite arborizations, axons fasciculation and cell bodies alignment are disturbed. **E)** Quantification of the VNC length in μm (average and standard deviation) for each condition above. Statistically significant differences in length (p < 0.0001) were detected between WT and (Hep^CA^ 10-16) and (Hep^CA^ 10-13) embryos. **F)** Kymographs (bright field) displaying muscle profiles at 25 hours AEL (100 sec) for the left (L) and right (R) sides of embryos expressing Hep^CA^ in RN2+ cells at early (10-13) or late (15-16) developmental stages. Anterior (A) and posterior (P). Embryos overexpressing *Hep*^*CA*^ at early stages in RN2+ neurons never managed to coordinate muscle movements but those overexpressing *Hep*^*CA*^ late, developed bilaterally symmetrical peristalsis (red arrows) and hatched in all occasions (**Movie S6**).

In conclusion, the condensation of the VNC and the coordination of the peristaltic movements leading to larval hatching are both dependent on precise levels of JNK activity in pioneer neurons at the beginning of CNS morphogenesis. Early, JNK modulates the establishment of the pioneer landscape and alterations in its activity induce aberrations in the axonal scaffold, in cell bodies positioning, and in the coordination of muscles activity. Late, JNK impinges upon the final refinement of the allocation of neurons and axonal tracts, but it is unrelated to motor control (**Figure 6H**).

## DISCUSSION

### JNK signaling coordinates motor activities

In the *Drosophila* embryo, muscles become active very early and isolated twitches and unilateral waves of contraction occur prior to the generation of propagated action potentials ^29^. This stochasticity eventually evolves towards organized coordinated muscle contractions that will promote the hatching of newborn larvae. We asked if pioneer neurons and the architectural organization of the VNC have any functional significance in this context. Our data strongly advocate for a close relationship between the acquisition of VNC architectural robustness and the rhythmic coordination of motor functions. Mutant embryos of different *puc* alleles and those overexpressing Hep^CA^ in pioneer neurons show severe structural defects (Karkali et al, co-submitted) and also slower and uncoordinated muscle twitching; they never hatch.

The failure of motor coordination could be explained as an indirect result of firing defects ^30^, associated to the aberrant dendritic arborizations of pioneer motoneurons observed when altering the level of JNK activity. Dendrites carry the vast majority of synaptic inputs and their morphology and properties are essential for propagation of postsynaptic potentials ^31^. Indeed, JNK signaling has been previously shown to promote dendrite pruning in *Drosophila* sensory neurons ^16^. Further, at the larval neuromuscular junction (NMJ), JNK signaling affects bouton number and alters the amplitude of excitatory junction potentials (EJP) and spontaneous responses. The sustained postembryonic JNK activation triggers synaptic growth, while the strength of the synapses is reduced ^8, 14^. Last, it has been suggested that JNKs might also be involved in developmental neuroplasticity as JNKs are activated by environmental stimulation and by electroconvulsive seizures in mice models ^20^.

A remarkable consequence of increasing JNK activity in pioneer motoneurons is the hyperpolarization of their membranes (**Figure 6**) and it is well known that the membrane potential can regulate cell differentiation as well as cortical tissue arrangement, shape and size. Conceivably, in RN2 neurons, synaptic integration may be affected by changes in dendritic arborization and morphology associated to high JNK activity. This in turn may affect the frequency or capability of firing action potentials. Indeed, it is known that hyperpolarizing conditions lead to a reduction of the number of connections in immature neurons *in vitro* ^32^. Along these arguments, the hyperpolarization of RN2 cells after Hep^CA^ overexpression may in part account for their anomalous dendritic arborization. Less probable would be a direct effect in the implementation of motor coordination. If this were the case, the hyperpolarization of RN2 cells should be linked to JNK early functions which, considering that action potentials generate at late embryonic stages ^29^, is unlikely. Our results thus suggest that muscle coordination failure will be primarily associated to the structural defects of the VNC after early hyperactivation of the JNK pathway in pioneer neurons (**Figure 7**).

**Figure 7.**
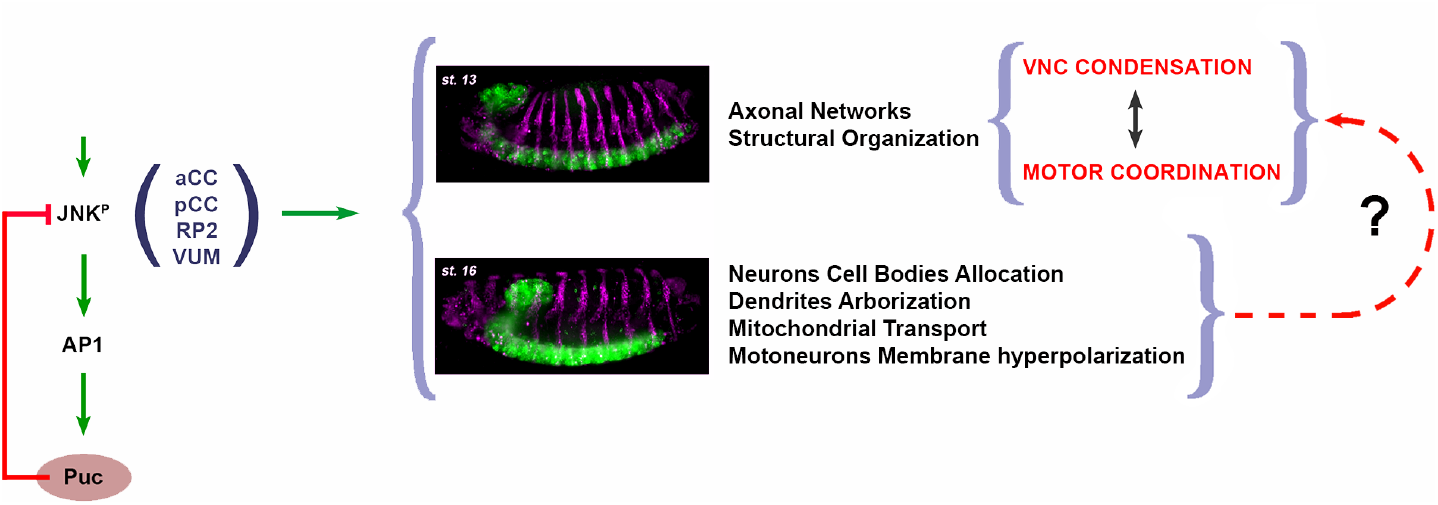
The early activity of the JNK pioneer neurons affects the structural organization of axonal networks and motor activity coordination. Precise JNK activity levels in early-specified neurons (at least aCC, pCC, RP2 and VUMs) are regulated by a negative feedback loop mediated by Puc. Excessive JNK activity in *puc* mutants or upon overexpressing HepCA in aCC, pCC, RP2 and/or VUMs leads to autonomous defects in axonal paths and dendrites numbers and shape, mitochondrial axonal transport and hyperpolarization of motoneurons membranes. Further, the architectural robustness of the VNC is affected, its condensation prevented and the coordination of embryonic motor activities and peristaltic movements inhibited. The incomplete condensation of the VNC may result in a failure in wiring optimization, partly responsible of motor uncoordination.

### JNK and the Cell Biology and Physiology of pioneer neurons

It has been reported that the JNK cascade in the CNS regulates microtubule dynamics by phosphorylating both, microtubule-associated proteins, including Tau ^33^, or the kinesin heavy chain (Khc), which induces the kinesin motor to release from microtubules ^11^. In the *Drosophila* VNC, Tau together with microtubule-binding spectraplakins modulates microtubule stability, which leads to alterations of JNK signaling activity, consequently affecting kinesin-mediated axonal vesicle transport ^13^. In our hands, however, interfering in RN2 pioneer motoneurons JNK activity does not significantly affect synaptic vesicles transport (**Figure S2**), although it inhibits mitochondrial motility (**Figure 3**), an effect previously observed in third instar larvae, where the JNK pathway mediates the inhibition of mitochondrial anterograde transport by oxidative stress ^12^. These data suggest that the influence of JNK signaling in conveying vesicles movements is probably cell and developmental stage specific and, significantly, indicate that synaptic vesicles axonal transport in pioneer neurons is irrelevant for embryos motor coordination. In addition, although it is known that JNK activity perturbs neuronal migration during cortical development in mice ^18, 34^, we have not observed any relevant drift of pioneer neurons when their level of JNK activity is altered.

While altering JNK signaling affects the cell biology (axonal transport) and physiology (hyperpolarization) of pioneer neurons, we do not know if it does so through its well-established role regulating gene expression, phosphorylating substrates such as Jun or histone modifiers, or, alternatively, acts via the phosphorylation of cytoplasmic targets. The fact that, in different models, the JNK pathway mediates microtubule destabilization leading to a transport roadblock by phosphorylating cytoplasmic substrates, including microtubule-associate proteins ^33, 35^, strongly supports this last possibility.

### Wiring Optimization

It is well-established that spontaneous activities that persistently stimulate postsynaptic cells result in long-term synapse potentiation ^36^. These activities propagate efficiently between nearby cells, ensuring connections are strengthened, whereas those from distant ones are lost ^37^. Bringing neurons in proximity improves functional interconnectedness and improves wiring optimization ^2^. We suggest that these classic principles are followed during the coordination of motor activities in the *Drosophila* embryo. Several arguments support this premise. First, both VNC condensation and motor coordination require a precise level of JNK activity at early developmental stages, enabling activity of pioneer neurons (**Figure 6**). Second, either activation or ablation of the EL interneurons disrupt bilateral muscle contractions in the larvae ^38^. Remarkably, these ELs make presynaptic contacts with the RP2 motoneurons, which we found critical for both, the architectural organization of the VNC (Karkali et al, co-submitted) and for motor coordination (**Figures 2** and **6**). Third, ELs, and the dorsal projecting motoneurons they innervate, are nearly all *puc*-expressing neurons (Karkali et al, co-submitted).

We foresee that the principles of wiring optimization may also apply at a postembryonic level. As larvae develop, their body wall area grows by two orders of magnitude ^39^. To accommodate such a change, the mechanosensory neurons must extend their dendrites and increase their receptive field ^40^, while motoneurons add more synapses to adapt to larger muscles ^39, 41^. During scaling, synaptic contacts between nearby neurons are sustained ^42^. In this scenario, a post-synaptic neuron can only connect with nearby specific pre-synaptic sites. The propensity to form stable local functional synapses appears to operate across development.

Here we uncovered an intimate causal link between the temporal control of the activation of the JNK pathway in pioneer neurons, the structural organization of axonal networks, VNC condensation, and embryonic motor coordination (**Figure 7**). Correlative geometrical and mechanical studies and further cell type-specific genetic interference analyses will be necessary to dissect the precise mechanisms underlying this process. Yet, wiring optimization across different circuits, by enhancing partners’ proximity and topographical robustness, appears to be an essential character associated with the morphogenesis of the nervous system.

## METHODS

### Drosophila strains

The following stocks were used:

*w1118*; *puc*^*E69*^*LacZ/TM3, twi-GFP* ^43^

*w1118*; *P{w[+mC]=UAS-Hep*.*Act}2* (BDSC #9306)

*w1118*; *P{w[+mC]=eve-Gal4*.*RN2}T, P{w[+mC]=UAS-mCD8::GFP*.*L}; P{w[+mC]=eve-Gal4*.*RN2}G, P{w[+mC]=UAS-mCD8::GFP*.*L}* (Dr. Irene Miguel-Aliaga)

*w1118 P{w[+mW*.*hs]=GawB}MzVUM; P{y[+t7*.*7]w[+mC] = 10xUAS-IVS-mCD8::GFP} attP40* (Dr. Irene Miguel-Aliaga)

*w1118*; *G203:UAS-TNT; ZCL2144* ^7^ (Dr. Matthias Landgraf)

*w1118*;; *ZCL2144* ^7^ (Dr. Matthias Landgraf)

*w1118; P{UAS-syt*.*eGFP}2* (BDSC #6925)

*w[1118]; P{w[+mC]=UAS-mitoGFP*.*AP}2 / CyO* (BDSC #8442)

In all cases, unless otherwise stated, embryos of the *w1118* strain served as controls.

### Genetics

All crosses were performed at room temperature and after 48 hours were shifted to different temperatures as the individual experiments required.

For temperature shifts, 2 hours embryo collections were done at room temperature in agar plates and aged for a further 2 hours. Then, they were transferred, either to a 29ºC incubator for 4 hours and then to an 18ºC incubator for 12 additional hours [Hep^CA^ (10-13)] or to an 18ºC incubator for 12 hours and then to a 29ºC incubator for 4 hours [Hep^CA^ (15-16)]. These timing regimes were calculated to enhance Hep^CA^ expression during the early pioneering stages or during the final cell allocation and axonal refinement stages of VNC development. After the temperature shifts the embryos were processed for immunocytochemistry or for live imaging.

### Immunohistochemistry

Immunostaining of flat-prepped stage 16 *Drosophila* embryos was performed using the primary antibodies: mouse anti-Fas 2 (1:100, clone 1D4, DHSB), and rabbit anti-GFP tag polyclonal (1:600, Thermo Fisher Scientific, #A11122).

The secondary antibodies used for detection were: Goat anti-Rabbit IgG (H+L), Alexa Fluor 488 conjugate (A-11008) and Goat anti-Mouse IgG (H+L) Alexa Fluor 555 conjugate (A-21422). All secondary antibodies were used in a dilution of 1:600 and were from Invitrogen.

### Sample preparations for immunodetection and image acquisition

*Drosophila* embryo dissections for generating flat preparations were performed according to ^25^. Briefly, flies maintained in apple juice-agar plates at 25ºC were synchronized by repetitive changes of the juice-agar plate, with a time interval of 2 hours. All embryos laid within this time window were aged at different temperatures following experimental requirements until reaching mid-stage 16 (3-part gut stage). At this point embryos were dechorionated with bleach for 1 min, poured into a mesh and rinsed extensively with water. For dissection, embryos were transferred with forceps on the surface of a small piece of double-sided tape, adhered on one of the sides of a poly-L-Lysine coated coverslip. After orienting the embryos dorsal side up and posterior end towards the center of the coverslip, the coverslip was flooded with saline (0.075 M Phosphate Buffer, pH 7.2). Using a pulled glass needle the embryos were manually de-vitelinized and dragged to the center of the coverslip, where they were attached to the coated glass with their ventral side down. An incision on the dorsal side of the embryo was performed using the glass needle from the anterior to the posterior end of the embryo. The gut was removed by mouth suction and a blowing stream of saline was used to flatten their lateral epidermis. Tissue fixation was done with 3.7 % formaldehyde in saline for 10 minutes at room temperature. After this point standard immunostaining procedures were followed.

Image acquisition was performed on a Zeiss LSM 700 inverted confocal microscope, using a 40X oil objective lens (NA 1.3). Z-stacks, spanning the whole VNC thickness, were acquired sectioning with a step size of 1 μm. Image processing was performed with Fiji ^44^.

### Live imaging

Dechorionated stage 14 embryos were glued ventral side down on a MatTek glass bottom dish and they were covered with S100 Halocarbon Oil (Merck) to avoid desiccation. Image acquisition was performed on a Zeiss LSM 700 inverted confocal microscope, using a 25X oil immersion lens (N.A 0.8, Imm Korr DIC M27). Z-stacks spanning the whole VNC thickness, with a 2 μm step size, were acquired every 6 minutes for a total of 8 hours. Processing of time-lapse data was done with Fiji ^44^.

### Image analysis

Image analysis and quantification of fluorescence intensity were performed using Fiji ^44^. In immunostainings where quantification of the fluorescent intensity was required, extra care was taken so that the same antibody batch was used for staining embryos of different genotypes, while identical confocal acquisition settings were applied.

### Statistical analysis

All statistical analyses were performed using GraphPad Software (GraphPad Software Inc., La Jolla, CA, USA). In all cases one sample t test was performed and probability values p<0.05 were considered as significant.

### Muscle contraction analysis

Stage 17 embryos carrying the G203, ZCL2144 or both GFP muscle markers were dechorionated and mounted in halocarbon oil (3:1 mix of S700 and S27) following standard procedures. Muscle contractions were monitored at the embryo’s middle plane, using a 25X oil immersion lens (N.A 0.8, Imm Korr DIC M27) in a Zeiss LSM780 confocal microscope. For each acquisition 1000 frames were obtained with a scanning speed of 200 msec per frame with a time interval between acquisition cycles of 15 minutes. Muscle activity was monitored up to embryo’s hatching or for a total of 12 hours at room temperature. For the experiments where Hep^CA^ over-expression in RN2 and MzVum neurons was required, both control and Hep^CA^ overexpressing embryos were filmed at 29°C for equal time periods. For temperature-shift analyses of animals expressing Hep^CA^ in RN2 neurons, embryos of equivalent pre-hatching age were recorded at 200 msec per frame for 500 frames. Post-imaging, movies were processed with Fiji ^44^ and Kymographs were generated using the Fiji plugin KymoResliceWide.

### Axonal Trafficking

To examine axonal trafficking, stage 17 embryos expressing UAS-Syt-GFP were dechorionated and mounted laterally in halocarbon oil (3:1 mix of S700 and S27) for live imaging. Synaptic vesicle transport was monitored in the medial part of the Inter-Segmental Nerve (ISN) using a 40X oil immersion lens (N.A 1.3, Oil DIC M27) in a Zeiss LSM780 confocal microscope. A constant frame size of 512×100 pixels, with an XY zoom of 6 and a recording line spacing of 2 were routinely employed. Single plane movies were acquired with a scanning speed of 1.11 sec, with 1 sec interval for a total of 60 cycles. All movies were processed with Fiji and Kymographs were generated using the Fiji plugin KymoResliceWide.

The Mean Velocity and the Lifetime parameters for each particle were calculated from the Kymographs as follows. The particles trajectories were traced and saved as independent ROIs. The XY coordinates for each ROI were then extracted using the getSelectionCoordinates function in Fiji of the custom made “Macro coordinates.ijm” shown below:

~~~
run (“Fit Spline”, “straighten”);
getSelectionCoordinates (x, y);
 x2=0; y2=0; distance=0;
 for (i=0; i<x.length; i++) {
 if (i>0) {
    dx = x[i] - x[i-1];
    dy = y[i] - y[i-1];
    distance = sqrt(dx*dx+dy*dy);
}
 print (i, x[i], y[i], distance);
}
~~~

These XY coordinates represent the particles displacement in both space (X values) and time (Y values), thus permitting the calculation of Mean Velocity and Lifetime.

### Analysis of Axonal and dendrite morphology by Segmentation

To describe the axonal and dendrite morphology of the RN2+ neurons, we analyzed images of flat-prepped embryos immunostained against GFP and Fasciclin 2 using the Simple Neurite Tracer Fiji plugin ^45^. Image segmentation by this tool allows a semi-automated tracing of axons and dendrites in 3D, providing measurements of total axonal and dendrite length, as well as of dendrite number directly from skeletonized images.

### Embryo whole-cell patch-clamp recordings

Dissections and recordings were performed as described ^29^ at room temperature (20-22°C). Late stage 17 *Drosophila* embryos were dissected in external saline (in mM: 135 NaCl, 5 KCl, 4 MgCl2·6H2O, 2 CaCl2·2H2O, 5 N-Tris[hydroxymethyl]methyl-2-aminoethanesulfonic acid, and 36 sucrose, pH 7.15). The CNS was exposed through a dorsal-longitudinal cut of the dechorionated embryonic body wall musculature. Embryonic cuticle was secured to a Sylgard (Dow-Corning, Midland, Michigan, USA)-coated cover slip using tissue glue (GLUture; WPI, Hitchin, UK) and larval gut and fat body removed. The glia surrounding the CNS was partially removed using protease (1% type XIV; Sigma, Dorset, UK) contained in a wide-bore (15 μm) patch pipette. Whole cell recordings were carried out using borosilicate glass electrodes (GC100TF-10; Harvard Apparatus, Edenbridge, UK), fire-polished to resistances of between 16-20 MΩ. Cell identity was confirmed through RN2-GAL4 driven expression of UAS-CD8-GFP. Recordings were made using a MultiClamp 700B amplifier. Endogenous excitatory synaptic cholinergic input to the aCC/RP2 motoneurons were recorded in voltage clamp and current clamp sampled at 20 kHz and low pass filtered at 10 kHz, using pClamp 10.6 (Molecular Devices, Sunnyvale, CA). Only neurons with an input resistance of ≥ 1GΩ were accepted for analysis.

## ACKNOWLEDGEMENTS

We are extremely grateful to all colleagues that have provided us materials, training and guidance throughout this work as well as for commenting on the manuscript. Amongst them we like to specially thank Christian Klambt, Claude Desplan, Matthias Landgraf, Chris Doe and Brian McCabe. We also thank our colleagues at the IBMB and the BSRC “Alexander Fleming” for their encouragement and support. The Martín-Blanco laboratory was supported by funds from Ministry of Innovation and Science (BFU2014-57019-P and BFU2017-82876-P) and Fundación Ramón Areces. RAB was supported by a BBSRC BB/L027690/1 grant and SWV by a BBSRC CASE studentship. Work on this project benefited from the Manchester Fly Facility with funds from University and the Wellcome Trust (087742/Z/08/Z).

## SUPPLEMENTARY FIGURES

**Figure S1.**
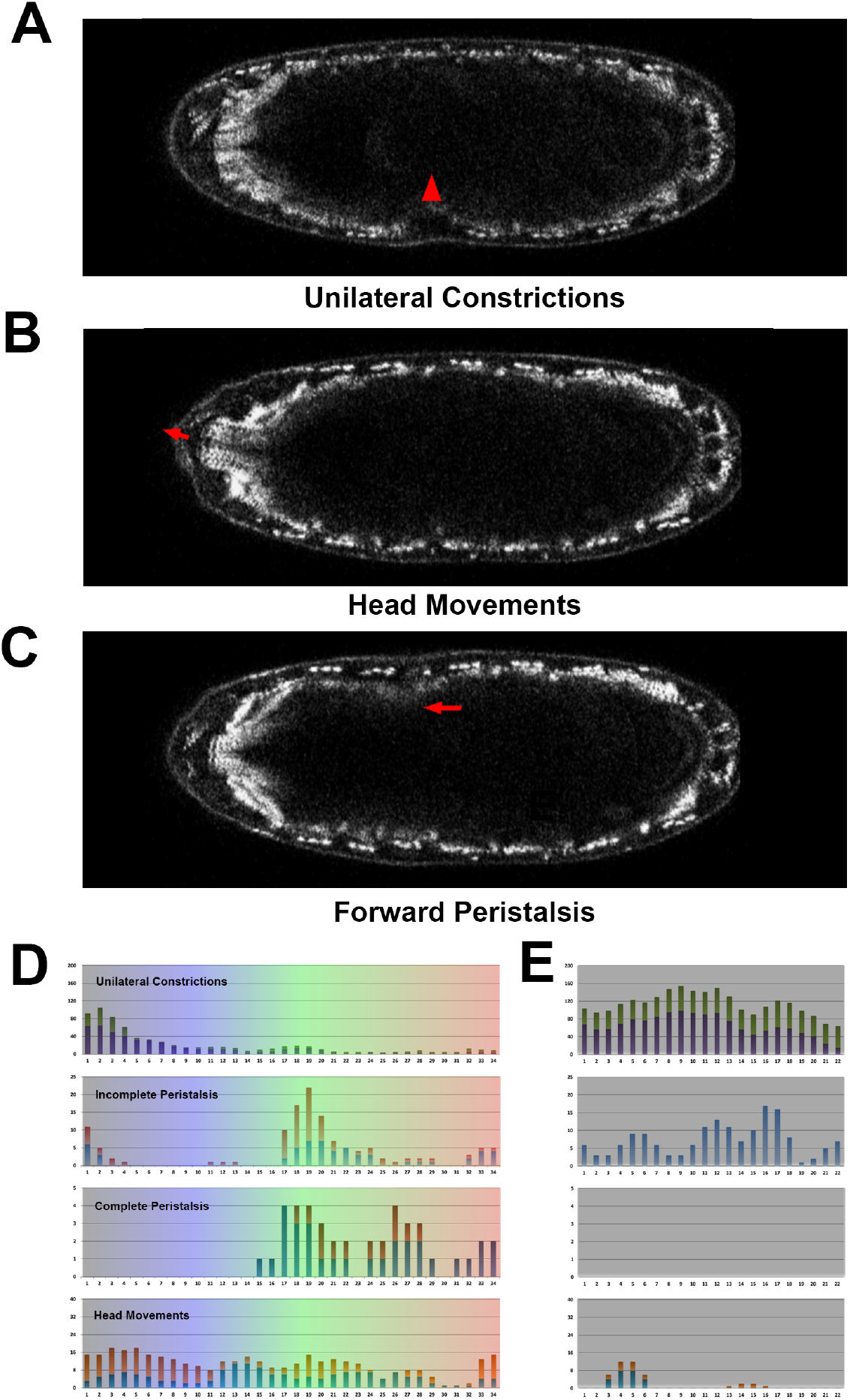
Types of embryonic muscle movements. Snapshots from **Movie S1** displaying mid-plane images of a stage 17 embryo carrying two GFP protein traps expressed at muscle Z-lines (*w; G203, ZCL2144*). Three main types of movement were observed during the muscle coordination process. Red arrows point to the contracting muscle units. Anterior is to the left. Scale bar is 50 μm. **A)** Unilateral muscle contractions. They can occur at any embryo side and in different segments. **B)** Head turning and mouth hook pinching. **C)** Peristalsis. They can be incomplete or complete and develop forward (in the image) or backward displacements. **D)** Histograms showing the progression of the muscle coordination process in wild type embryos. Over-imposed colors follow the 5 stages of motor coordination maturation described in **C**. Muscle movements were distributed in four classes: Unilateral contractions [sum of the contractions occurring at the right (green) and the left (purple) side of the embryo]; Incomplete peristalsis [sum of the incomplete forward (blue) and incomplete backward (red) peristaltic attempts]; Complete peristalsis [sum of the full forward (blue) and backward (red) peristalsis]; and Head movements [sum of head turning (red) and mouth hook pinching events (blue)]. **E)** Histograms equivalent to **D** showing the unsuccessful muscle coordination of *puc*^*E69*^ embryos.

**Figure S2.**
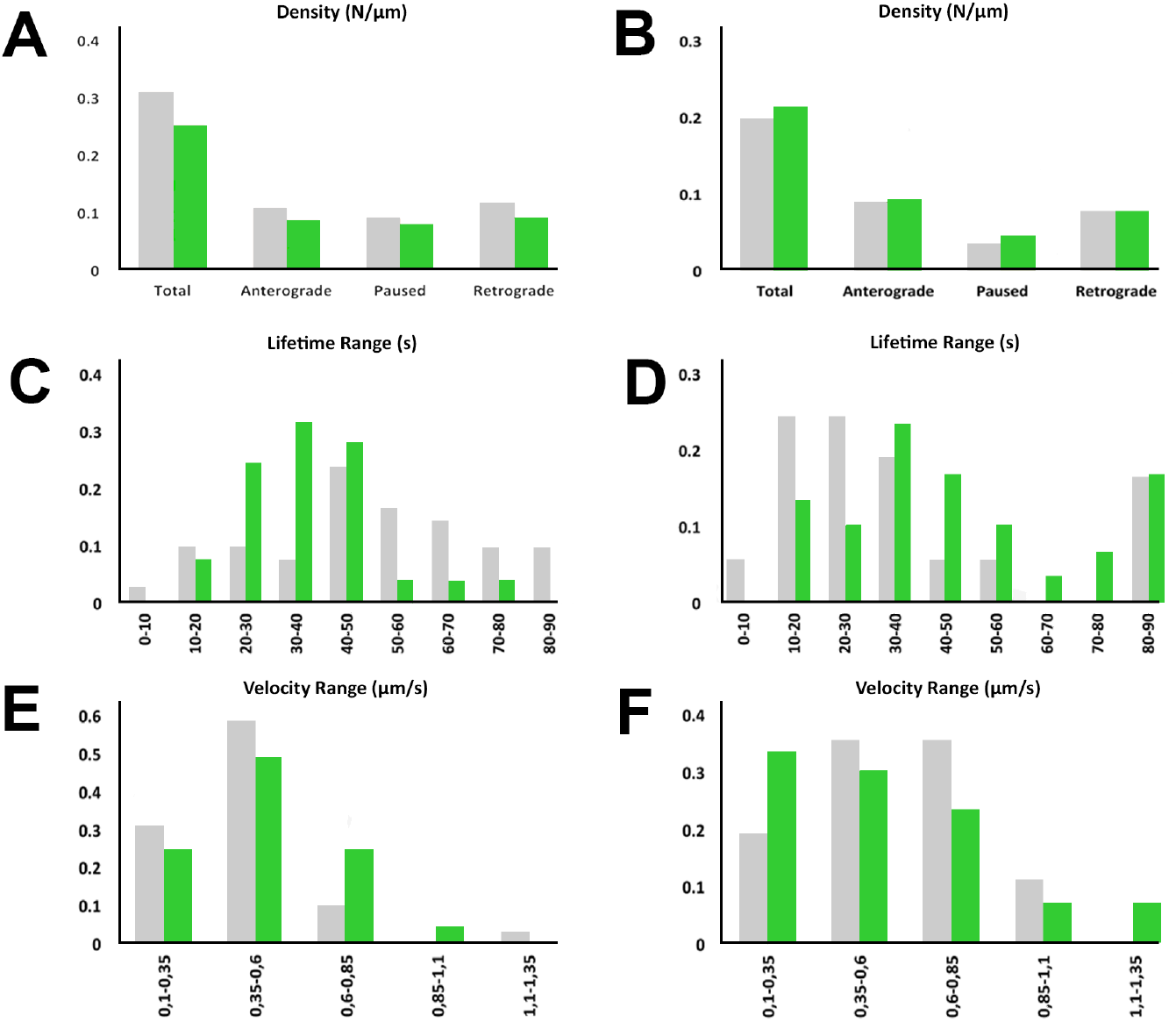
Synaptotagmin transport is unaffected by JNK hyper-activation in aCC/RP2 and VUM motoneurons. A and B) Histograms showing synaptotagmin vesicles density and directional motility in control (grey) versus Hep^CA^ expressing (green) aCC/ RP2 (A) and VUM motoneurons (B). No statistically significant differences were observed between control and JNK over-active neurons. C and D) Lifetime range distribution of motile synaptotagmin vesicles in aCC/RP2 (C) and VUM (D) motoneurons. No differences were observed in lifetime between control (grey) and Hep^CA^ expressing (green) embryos. E and F) Mean velocity range distribution comparison of motile synaptotagmin vesicles in aCC/RP2 (E) and VUM (F) motoneurons between control (grey) and JNK gain-of function (green) conditions. For both sets of motoneurons the distribution was unaltered.

## SUPPLEMENTARY MOVIES LEGENDS

**Movie S1. Embryonic motor patterns**

Representative mid-plane movies of a stage 17 embryo carrying two GFP protein traps expressed at muscle Z-lines (*w; G203; ZCL2144*). **A)** Forward peristalsis. **B)** Backward Peristalsis. **C)** Left twitch. **D)** Right twitch. **E)** Head tumble. **F)** Mandible pinch. Scale bar is 50 μm. Anterior is to the left.

**Movie S2. Time course of motor maturation**

Representative mid-plane movies (250 time points per stage) of an embryo carrying two GFP protein traps expressed at muscle Z-lines (*w; G203; ZCL2144*). From top to bottom stages **A** (Unilateral twitches and incomplete peristalsis), **B** (Resting period), **C** (Complete peristalsis), **D** (Late resting period) and **E** (Hatching movements). Color coded as in **Figure 6**. Scale bar is 50 μm. Anterior is to the left.

**Movie S3. Motor activity of *puc***^***E69***^ **embryos**

Representative mid-plane movie (1000 time points) of a late stage 17 *puc*^*E69*^ embryo carrying two GFP protein traps expressed at muscle Z-lines (*w; G203; ZCL2144*). Scale bar is 50 μm. Anterior is to the left.

**Movie S4. Synaptic vesicles axonal transport in RN2 neurons**

Stage 16 embryos expressing Synaptotagmin-GFP under the control of the RN2-Gal4 in the absence **(A)** or presence **(B)** of Hep^CA^. Left panels present representative single plane movies (Fire Luts) with 1 sec interval for a total of 60 cycles. Right panels display in different colors the traces of the different particles analyzed. Proximal is to the left. Scale bar is 25 μm.

**Movie S5. Axonal and dendritic landscape of pioneer neurons upon JNK hyper-activation**

Animated 3D reconstruction of the axonal and dendritic pattern of stage 16 embryos expressing mCD8-GFP (**A**) and mCD8-GFP and Hep^CA^ (**B**) under the control of the RN2-Gal4 line. Traces of the dendrites (purple) are over-imposed. Scale bar is 10 μm.

**Movie S6. Motor coordination responses to early and late JNK hyperactivation**

Representative mid-plane movies of stage 17 embryos expressing Hep^CA^ in RN2+ cells at early (13 - top) or late (16 - bottom) developmental stages. Embryos over-expressing Hep^CA^ at early stages never coordinate muscle movements but those overexpressing Hep^CA^ late, developed multiple bilaterally symmetrical peristalsis and hatched. Scale bar is 50 μm

